# Genotypic and functional characterization of fibroblasts derived from pressure sores

**DOI:** 10.64898/2026.05.21.726782

**Authors:** C Boyer, A Coste, E Tournier, B Chaput, B Sallerin, A Varin, S Gandolfi

## Abstract

**Introduction:** Pressure sores are a major health problem in people with spinal cord injury resulting in ischaemic tissue lesions caused by prolonged pressure against a bony surface. Conventional therapies are often defective and fundamental researches on the healing process of pressure sores must be enriched in order to understand any novel therapies that may be applied. We focalize on pressure sore’s fibroblasts as dermal fibroblasts perform a critic role in wound healing by populating the wound site to produce extracellular matrix. After characterizing morphological and the genetic profile of healthy fibroblasts and fibroblasts from pressure ulcers, we conducted an analysis of fibroblast proliferation, migration and myofibroblastic differentiation capacity.

**Materials and Methods:** after acquisition of dermal explants and fibroblasts culture, we conducted histological analysis, an evaluation of gene expression by RT-qPCR and an assessment of fibroblasts proliferation and migration capacity through IncuCyte. A study of the differentiation of fibroblasts into myofibroblasts through the detection of Alpha-Smooth Muscle Actin (α-SMA) expression by immunofluorescence was also conducted.

**Results:** histological analysis showed histological analysis showed dermal disorganization in pressure sore compared with health skin, differences in morphological aspects and density of fibroblasts.

Pressure sore fibroblasts express less genes coding for ECM proteins, metalloproteases, collagen III, Connective tissue growth factor (CTGF) and ACTA2 coding for α-SMA. Pathological fibroblasts appear to proliferate less quickly than healthy fibroblasts but no differences in migration capacity were found. After stimulation under TGF-β, pressure sore fibroblasts lose their ability to differentiate into myofibroblasts compared to healthy fibroblasts and this could be in relation with a less expression of ACTA2.

**Conclusion:** All of our results highlight a morphological, genetic and functional difference between healthy and pathological fibroblasts which have a modified phenotype, less effective for skin repair. This suggests that new therapies for chronic wounds must take into account the environment in which they are applied and that pathological cells do not necessarily respond to treatments in the same way as healthy cells. Our results are not statistically significant, although several trends emerge. This is explained by the heterogeneity of the patients’ medical history and requires repetition of the experiments.

## 1. Introduction

Pressure sores are ischaemic tissue lesions caused by prolonged pressure against a bony surface,when interstitial pressure exceeds 30 mmHg (arteriolar pressure) [1].

The resulting ischaemia affects tissues ranging from the skin to subcutaneous tissues, muscles and fascia simultaneously [2,3]. Ultimately, the ischaemia causes necrosis, leading to deep tissue ulceration. The development of skin necrosis in a pressure ulcer is therefore the result of tissue hypoxia and ischaemia-reperfusion injury.

The most widely used classification is that of the 2007 United States National Pressure Ulcer Advisory Panel (US-NPUAP), which was revised in 2016 and is the international benchmark [4-6]. Stage I pressure ulcers are erythematous or negative pressure areas. Clinically, stage I does not extend beyond the skin, although histological studies show that tissue damage extends beyond the skin barrier, with modifications in vessels in papillary dermis [7-9]. Stage II corresponds to a loss of cutaneous tissue and exposing the dermis. Stage III involves full-thickness damage to the skin barrier, exposing adipose tissue to varying degrees depending on the location of the pressure ulcer. If the pressure and aggravating factors persist, extensive necrosis defines a loss of substance that extends as far as the bone leading to septic complications (stage IV).

Pressure sores are a major problem in people with spinal cord injury. These patients suffer from sensory and/or motor disorders secondary to damage to their central nervous system, of encephalic or spinal cord origin.

The mechanism of injury may be sudden in onset, such as a stroke or direct trauma, or more progressive, as in multiple sclerosis or tumour compression. In case of spinal cord injury, there are numerous physiological and biomechanical changes.T hese changes affect both the spinal cord and the soft tissues below the level of injury.

Initially, vascular changes appear in the spinal cord:

- Vascular changes such as microhaemorrhages and vasospasms, at the origin of inflammation and changes in the permeability of the blood-brain barrier [10].
- The formation of free radicals, caused by cell oxidation, results in damage to the cell membrane, which also increases the local inflammatory reaction [11].
- Apoptosis, induced by cellular suffering. Apoptosis affects neurons, oligodendrocytes, microglia and astrocytes [12].

In a second stage, the inflammation generated leads to demyelination in the white matter of the spinal cord, while the grey matter dissolves. Scar tissue is produced in the spinal cord, acting as a barrier to axon regrowth and thus to neuronal electrical information. At the sub-lesional level, the diameter of muscle fibers decreases by a third. They become infiltrated with fatty tissue and fibrosis. The muscles atrophy, limiting the amount of fat padding on the skin in relation to the bone [13].

Ischaemic lesions are located in areas where stress is greatest. Sitting is the main cause of ischial lesions, while prolonged dorsal position is more likely to damage the sacrum, the trochanteric regions, as well as the heels, and malleolus during external rotation of the hip. In any case, the skin barrier is weakened by the pressure sore. Wounds are colonised by commensal germs, normally present in the physiological state of the skin. This microbiota plays a crucial role in the development of the wound. These bacteria maintain a balance by limiting the implantation of saprophytic, potentially pathogenic germs [14].

When the balance between commensal and saprophytic germs is unbalanced, pressure sores can lead to superinfection, either at the necrotic stage, before debridement, or afterwards when the underlying tissues are exposed [14].

If left uncontrolled or neglected, these wounds can lead to delayed healing and have serious repercussions on patients’ state of health, including septic shock, the risk of secondary carcinogenesis, limb amputation and even life-threatening conditions [13,15].

Conventional therapies are often insufficient and patients require debridement surgery and flap coverage [16]. Often these patients suffer from recurrences and relapses despite surgery because of their affection, that does not allow them adequate mobilization. In these circumstances the surgeon faces a therapeutic impasse because of the lack of tissue to mobilize and sometimes there is no longer any surgical solution. In this context, fundamental researches on the healing process of pressure sores must be enriched in order to understand any novel therapies that may be applied. We focalize on pressure sore’s fibroblasts as they are the cell population of the dermis, the first layer of skin that gives the skin its strength and solidity and forms the basis of the skin’s architecture and dermal fibroblasts perform a critic role in wound healing by populating the wound site to produce extracellular matrix (ECM) [17]. After characterizing morphological and the genetic profile of healthy fibroblasts and fibroblasts from pressure ulcers, we conducted an analysis of fibroblast proliferation, migration and myofibroblastic differentiation capacity.

## 2. Material and methods

### 2.1 Acquisition of human dermis explants from surgical specimens and fibroblasts culture

Surgical specimens were obtained from patients at the Department of Plastic and Reconstructive surgery of the Toulouse University Hospital. The study was conducted in accordance with the Declaration of Helsinki [18], and patients were informed about the research protocol and gave their free and informed consent (Collection N° DC-2020-4124). These were patients with chronic pressure sores for which debridement and flap coverage surgery was indicated and planned. Demographic characteristic of the population and risk factors for inadequate wound healing have been identified, such as obesity, diabetes, active smoking and recurrences. At the time of surgery, the plastic surgeon harvested a sample of the peri-lesional skin of the pressure sore (pressure sore sample - PS) and from the recut of the coverage flap, which is generally sacrificed at the end of the operation (healthy skin sample - H). Part of the periwound area was sent for anatomopathological examination.

Surgical specimens were kept in Phosphate-buffered saline (PBS) after collection, during the duration of transport. Under biosafety cabinet (BSC), skin specimens were washed in a PBS solution containing 2% ASP (Amphotericin, Streptomycin, Penicillin). The epidermis and hypodermis are mechanically removed and the dermis is preserved and cut into small explants (2 mm3) which are placed on petri dishes. The explants were left to air for 30 minutes and then cultured in α-MEM (α-Modified Eagle’s Medium) supplemented with 20% Fetal Bovine Serum (FBS), 1% ASP and 1% L-Glutamine (v/v). The medium was changed weekly until the fibroblasts migrated from the explant and colonized the petri dish (3 weeks) The fibroblasts were maintained at 37°C in a humid atmosphere containing 5% CO2. Figure 1 shows a schematic representation of the procedure.

**Fig.1:**
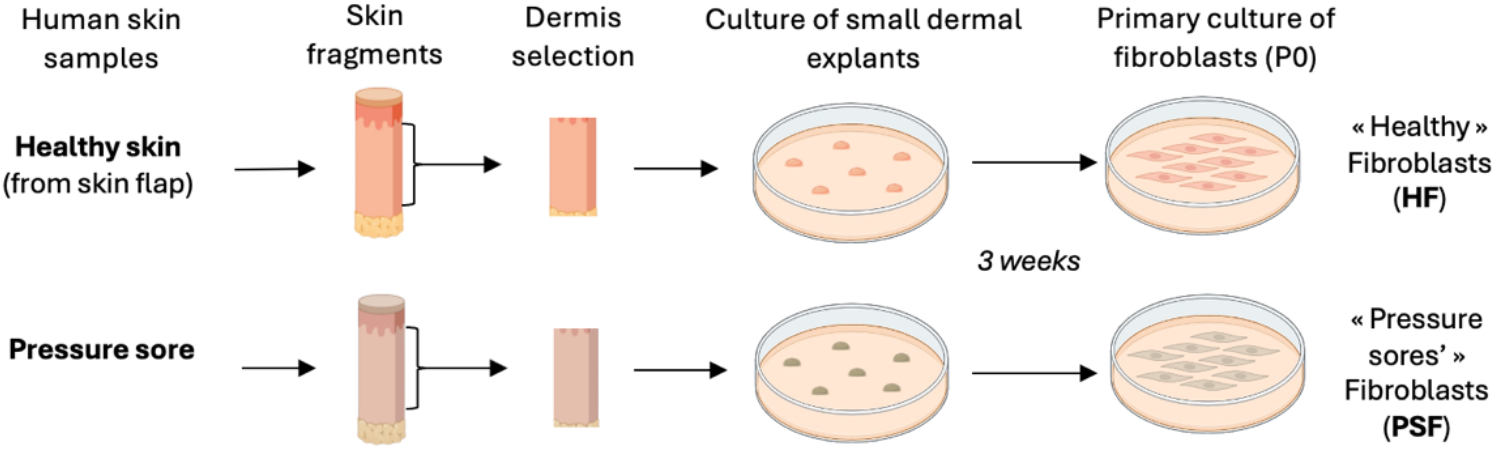
Schematic representation of the procedure.

When the fibroblasts were at 80% confluence, the pieces of dermis were removed from the petri dishes and the fibroblasts are detached using trypsin-0.05% EDTA (Gibco) and recultured in flasks at 2,000 cells/cm2 in α-MEM supplemented with 10% SVF, 1% ASP and 1% L-Glutamine (v/v). After one week in culture, the fibroblasts were detached and recultured at 2,000 cells/cm2. After a further week in culture, the fibroblasts were detached and frozen in 90% FBS + 10% cell freezing medium-DMSO (dimethyl sulfoxide). Fibroblasts between passages P1 and P3 were used for our tests (Fig.2).

**Fig.2:**
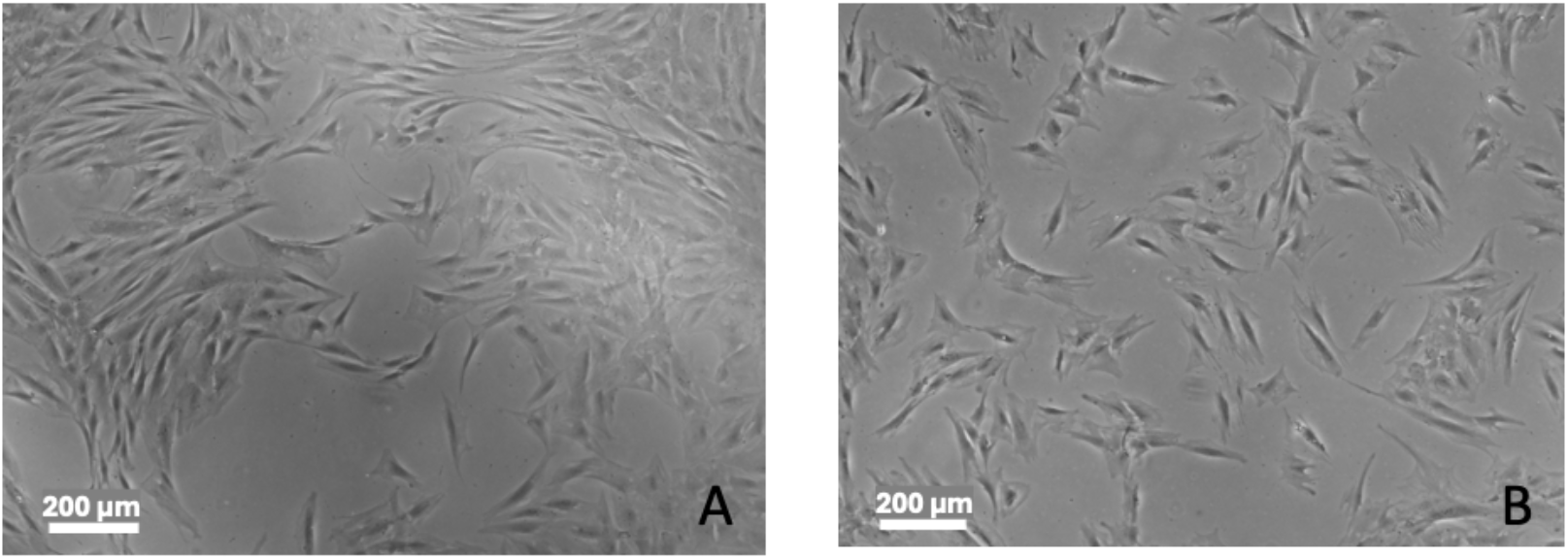
Representative photos of healthy fibroblasts (A) and fibroblasts from the eschar area (B) obtained after 2 passages in culture. Photographs are taken on an inverted microscope (Nikon) at 10X magnification. Scale bar: 200 μm.

### 2.2 Histological Analysis

The histological analysis of the surgical specimens was carried out in the anatomopathological laboratory of the university hospital center of Toulouse. All the pressure sores were examined to ensure that carcinomatous transformation was not overlooked. Semi-quantitative and semi-qualitative analysis was performed on 3 samples of fibroblasts from pressure sores and fibroblasts from healthy surgical specimens. Samples were fixed in formol solution, then treated with Heamatoxylin and Eosin (H&E) staining of paraffin sections.

### 2.3 Evaluation of gene expression in fibroblasts by RT-qPCR

Total RNA from HF (healthy fibroblasts) and PSF (pressure sores fibroblasts) was extracted using the “Quick RNA-Microprep” kit (Zymo Research) according to the manufacturer’s instructions. The quantity of RNA extracted was determined spectrophotometrically on the NanoDropTM 1000 (ThermoFisher Scientific). The total RNA is then reverse transcribed into cDNA using the High Capacity cDNA reverse transcription kit (Applied Biosystem) according to the supplier’s instructions.

qPCR was performed using SYBR Green reagent on the LightCycler® 480 II instrument (384-well format) (Roche Sequencing Solutions). The amplification conditions were as follows: 2min at 95°C (denaturation), then 60 cycles containing the following steps: 10s at 95°C (dehybridisation), 10s at 60°C (primer hybridisation) and 10s at 72°C (amplification). The amplification cycle is followed by the progressive denaturation of the amplification products (melting curves) by a 5s step from 65°C to 95°C. Serially diluted samples of pooled cDNA were used as external standards in each run for quantification. The primer sequences for each specific gene and the reference gene (GAPDH) used (Eurogentec) are detailed in Table 1.

**Table 1:**
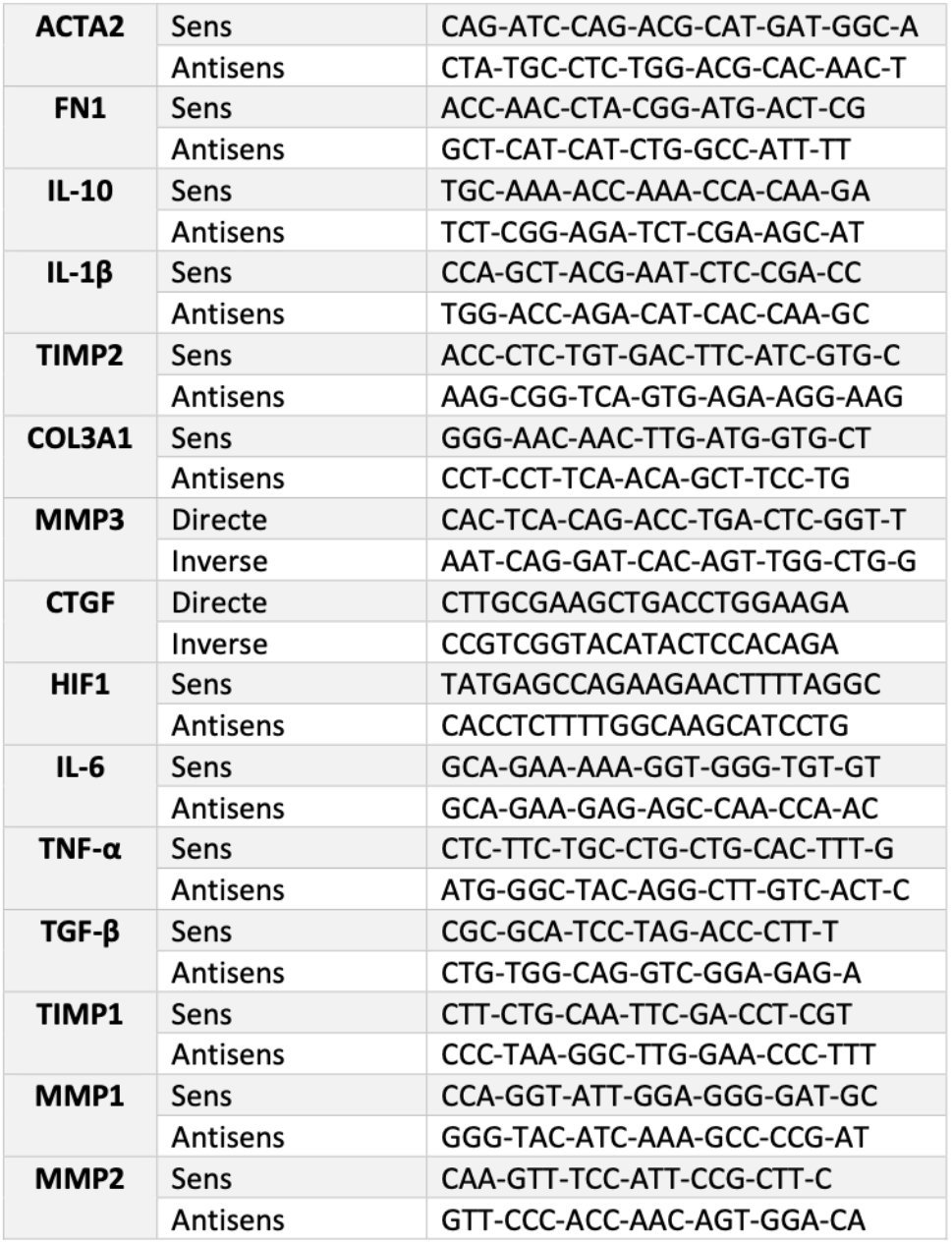
Summary table of primers used for RT-qPCR.

### 2.3 Assessment of fibroblast proliferation capacity

The different populations of fibroblasts were seeded at 5,000 cells per cm2 and were cultured in α-MEM + 10% SVF + 1% ASP. The plate was placed in the IncuCyte (Sartorius) and a image was taken every 6 hours for 7 days using a 10X objective. The analysis, carried out by the IncuCyte’ssoftware, is performed at all times and consists of establishing a confluence area recognising the cytoplasm of the cells. This area is used to monitor the confluence of cells over time and therefore to assess the proliferation of the different populations of fibroblasts. The proliferation curves presented are obtained as follows:

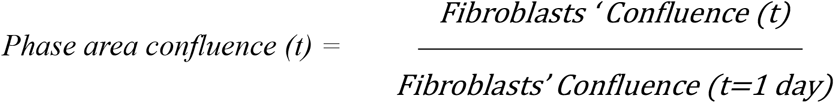

### 2.4 Assessment of fibroblast migration capacity

Human fibroblasts were seeded in a 96-well ImageLock plate (Sartorius) at 20,000 cells/well in α-MEM + 10% SVF + 1% ASP. After 4 h of adhesion, a wound was made in each well using the WoundMaker (Sartorius). After washing with PBS, the cells were also cultured in α-MEM + 10% SVF + 1% ASP (positive control) or α-MEM + 0.1% SVF + 1% ASP (negative control). The plate was placed in the IncuCyte and an image was taken every hour for 24 hours using a 10X objective. The analysis, carried out by the IncuCyte software, is performed at all times and is based on the calculation of the Relative Wound Density parameter. This is used to assess wound closure over time, and therefore fibroblast migration, and corresponds to the following ratio:

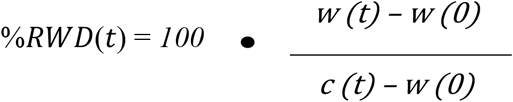

*w(t) density of the area at time t*

*c(t) density of the area occupied by the cells at time t*

### 2.5 Differentiation of fibroblasts into myofibroblasts: detection of Alpha-Smooth Muscle Actin (α-SMA) expression by immunofluorescence

Fibroblasts were cultured in α-MEM + 0.1% SVF + 1% ASP (negative control) 2 ng/mL TGF-β was added in culture condition. After 3 days of culture, the cells were washed with PBS and fixed with Paraformaldehyde (PFA) (0.37%) for 10 min at room temperature. After 3 washes with PBS, the cells were incubated for 30 min under agitation at room temperature in a PBS solution containing 2% NHS (Newborn Horse Serum, Thermofisher scientific) and 0.2% Triton, allowing saturation of the aspecific binding sites and permeabilisation of the membranes. Fibroblasts were then marked with a primary antibody against α-SMA (Mouse IgG, Zytomed) diluted 1:100 in PBS + 2% NHS + 0.2% Triton overnight at 4°C with agitation. After 3 washes in PBS + 0.2% Triton, the cells were incubated with an antibody coupled to an Alexa 488 fluorochrome that specifically recognizes the constant fragment of the primary mouse antibody (Goat anti-mouse IgG, Life Technologies, dilution 1/250 in PBS + 0.2% Triton).After 2 hours incubation at room temperature with agitation, the cells were washed with PBS and then incubated for 30 minutes at room temperature with a solution of 2-(4-Amidinophenyl)-6-indolecarbamidine (DAPI) staining solution (Sigma, 1/10,000 in PBS) and Alexa594-coupled phalloidin (Invitrogen, 1/400) with agitation, allowing labelling of nuclei and actin fibers respectively. After 2 washes with PBS, the cells were stored at 4°C in PBS.The markings obtained were analyzed using a high-throughput imaging system (Operetta, PerkinElmer) to quantify the fluorescence and determine the number of α-SMA-positive cells.

### 2.6 Statistics

The significance of the results obtained was assessed using Student’s t-tests and ANOVA performed on Prism 5 Software (GraphPad).

## Results

### Histological analysis

The histological sections examined revealed in all pressure sores significant changes showing an irregular, hyperplastic, hyperkeratotic epidermis with ulcerated areas replaced by a fibrin-leukocyte coating. The dermis exhibited a fibro scarring aspect and chronic inflammation in the architecture of the papillary and reticular dermis with the presence of focal necrosis, leucocytes infiltrates and an important fibrosis. No signs of malignancy or acute infections were found. The healthy skin showed no inflammatory changes and no changes in histological cutaneous architecture (Figure 3).

**Fig.3:**
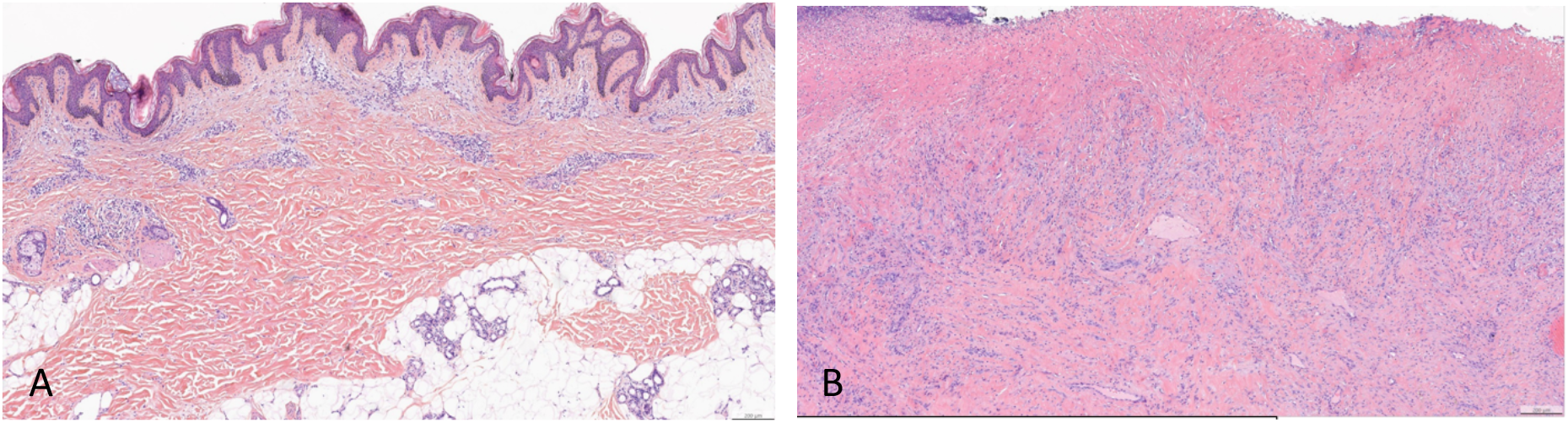
histological section of healthy skin (A) and pressure sore (B) of the same patient. Healthy skin keeps the normal architecture of cutaneous barrier (A). In pressure sores, the dermis exhibited a fibro scarring aspect and chronic inflammation (B) (scale 200 µm)

Pressure sores fibroblasts appeared enlarged and plump compared with healthy fibroblasts appearing thin and elongated (Figure 4).

**Fig.4:**
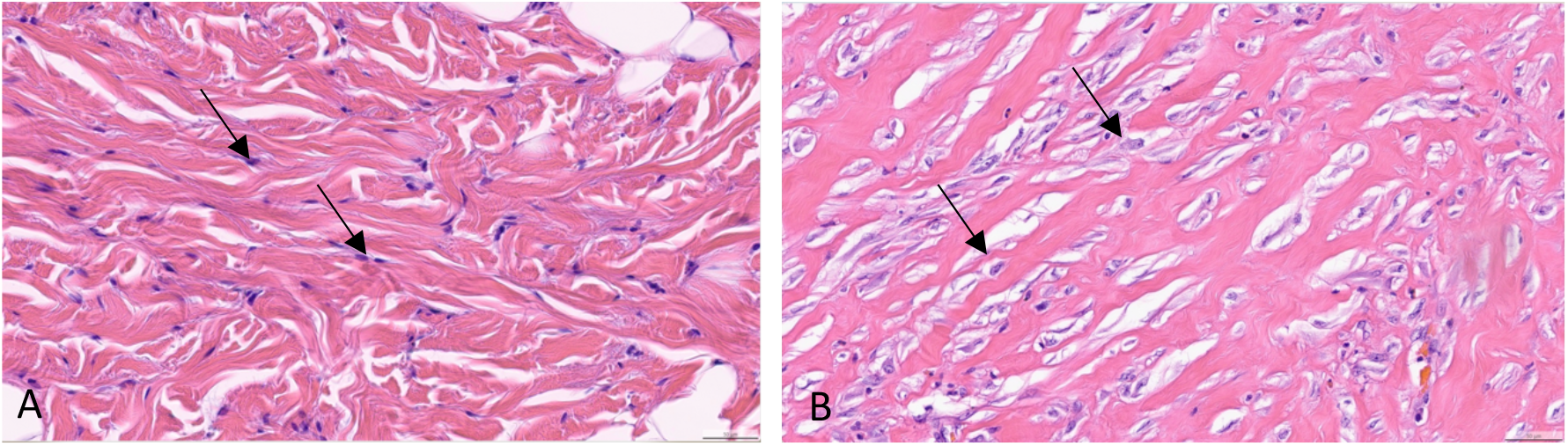
Morphological differences between healthy (A) and pathological fibroblasts (B) of the same patient. Healthy fibroblasts appear thin and elongated (A), pressures sores fibroblasts appear enlarged (B).

A lower density of fibroblasts in healthy samples compared with pathological sample was observed (1+ vs 3+ at x20 magnification).

### Gene expression

To compare the phenotype of the two types of fibroblasts, we studied the expression of genes encoding matrix molecules (COL3A1 and FN1), genes encoding proteins involved in extracellular matrix remodelling (TIMP2, MMP1, MMP2, MMP3), the ACTA2 gene encoding for α-SMA used as a marker of myofibroblast formation and genes encoding for citokines (IL-6, IL 10, ILb, TNFalfa, TGF-β) and growth factors (Connective tissue grow factor CTGF, and Hipoxia inducible-factor HIF-1).

Fibroblasts from the pressure sore (PSF) express fewer genes encoding Extracellular matrix (ECM) proteins such as COL3A1 (HF: 1.34 ± 0.11 vs PSF: 0.71 ± 0.31) and FN1 (HF: 1.42 ± 0.11 vs PSF: 1.07 ± 0.21) than healthy fibroblasts (HF) and showed decreased expression of genes encoding proteins involved in tissue degradation and remodelling such as MMP3 (HF: 1.01 ± 0.43 vs PSF: 0.63 ± 0.38) and TIMP2 (FNE: 1.20 ± 0.07 vs FE: 0.98 ± 0.18). Pathological fibroblasts also expressed less of the gene encoding CTGF (HF: 1.27 ± 0.12 vs PSF: 0.77 ± 0.26), a growth factor which stimulates cell proliferation, angiogenesis and ECM production by fibroblasts. In addition, healthy fibroblasts expressed more of the ACTA 2 gene, encoding the α-SMA protein, which is over-expressed in myofibroblasts (HF: 1.3 ± 0.47 vs PSF: 0.61 ± 0.12). However, pathological fibroblasts expressed more genes encoding pro-inflammatory cytokines such as IL-1β (FNE: 0.79 ± 0.38 vs FE: 1.56 ± 0.65) but also anti-inflammatory cytokines such as IL-10 (FNE: 1.19 ± 0.53 vs FE: 1.73 ± 0.74) (Fig. 5) Expression of IL-6, HIF-1, TGF-β and MMP1, MMP2, TIMP1, remained unchanged between the 2 fibroblast populations.

**Fig.5:**
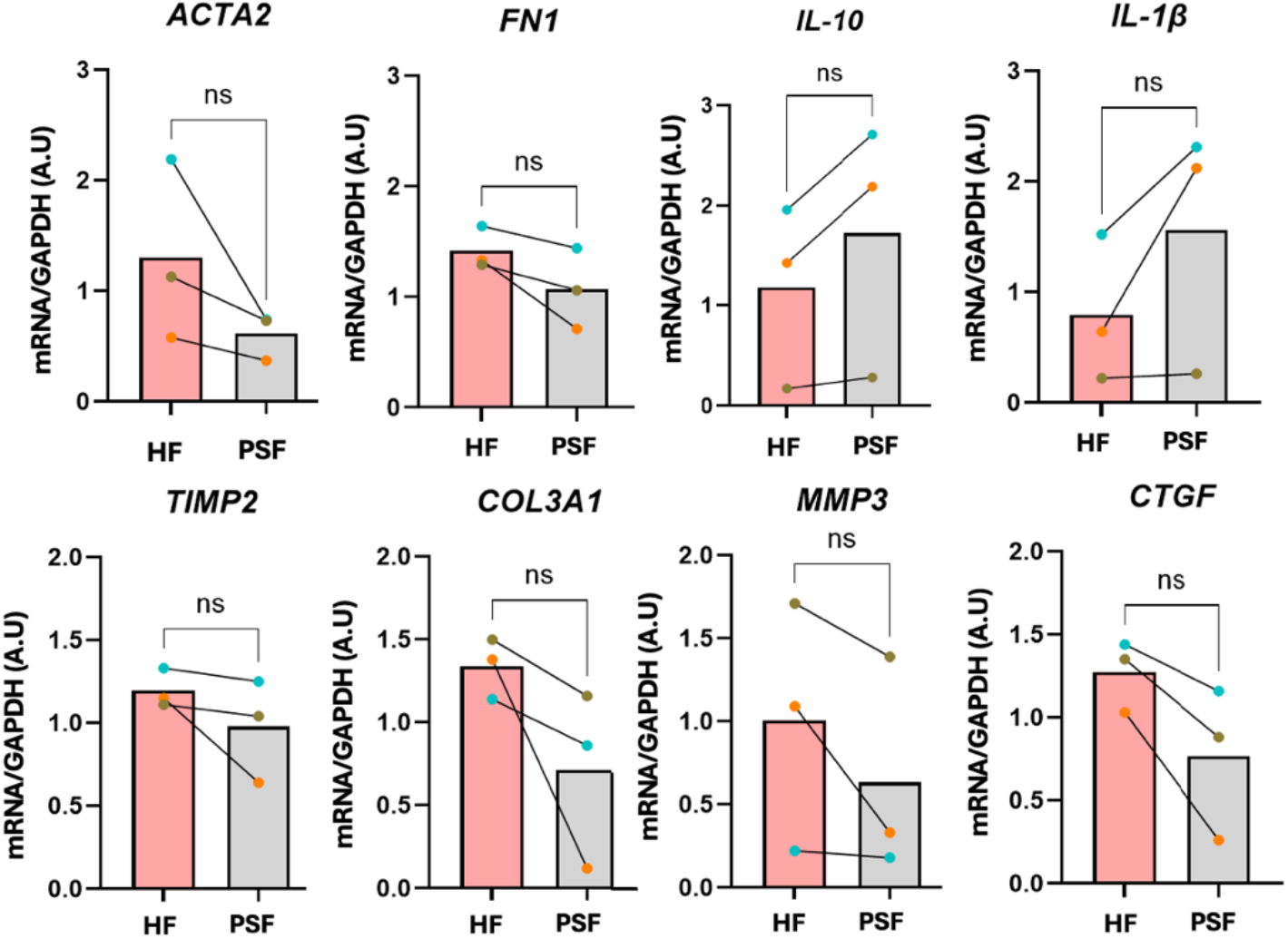
Basal gene expression of genes of interest from healthy fibroblasts (HF) and pressure sores fibroblasts (PSF) isolated from 3 different patients; (paired non-parametric t-test, ns: not significant). (GAPDH A.U: glyceraldehyde-3-phosphate dehydrogenase arbitrary unit).

### Fibroblasts’ proliferation and migration capacity

The basal proliferation capacity of healthy (HF) and pathological (PSF) fibroblasts was studied and monitored every 6 h for 7 days at the 10X objective using IncuCyte. Both fibroblast populations were cultured in basal medium (α-MEM supplemented with 10% SVF). PSF appeared to proliferate less than HF (Area under the curve AUC HS: 113.9 ± 16.31 and AUC PSF: 92.26 ± 22.62) (Fig.6 A,B) However, there was some heterogeneity between samples, with 1 sample showing no difference in proliferation between the 2 types of fibroblasts (Fig.6 B). No difference in migration capacity is observed under standard medium for each population fibroblasts (HF, PSF).

**Fig.6:**
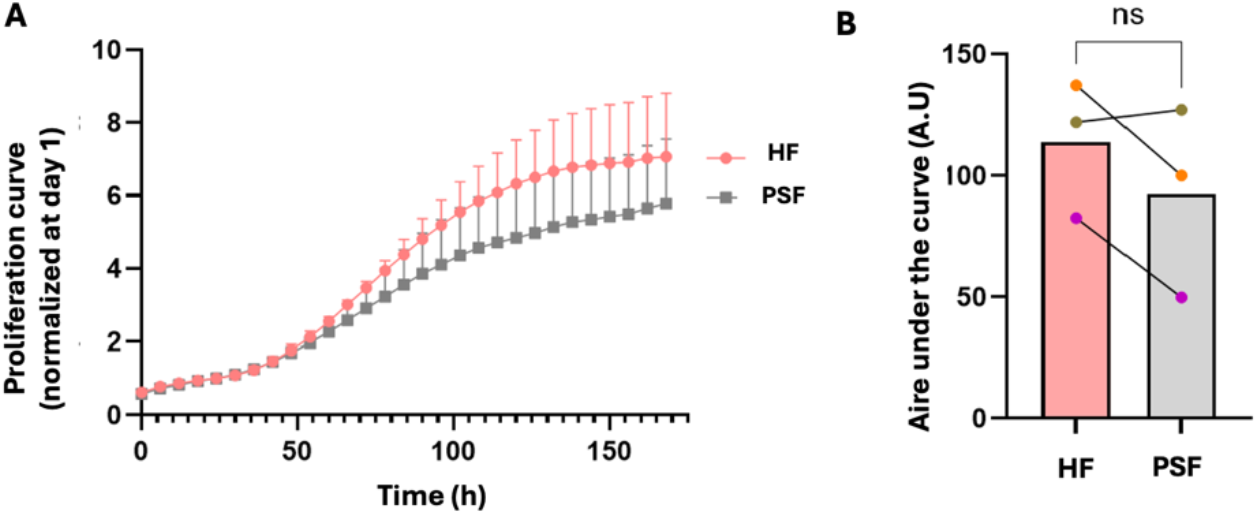
Basal proliferation capacity of healthy fibroblasts (HF) and pressure sore fibroblast (PSF). A: Graphical representation of HF and PSF proliferation over time. B: Mean areas under the proliferation curve (AUC), with each point representing a different patient sample. (n = 3 paired independent samples; non-parametric paired t-test (ns: not significant).

### Myofibroblast differentiation capacity

After stimulation under TGF-β, pressure sore fibroblasts (PFS) lose their ability to differentiate into myofibroblasts compared to healthy fibroblasts (HS) (HS: 21.61% ± 3.70 vs PFS: 3.56 ± 1.36% α-SMA+ cells/total cells) (Fig. 7 A-B)

**Fig.7:**
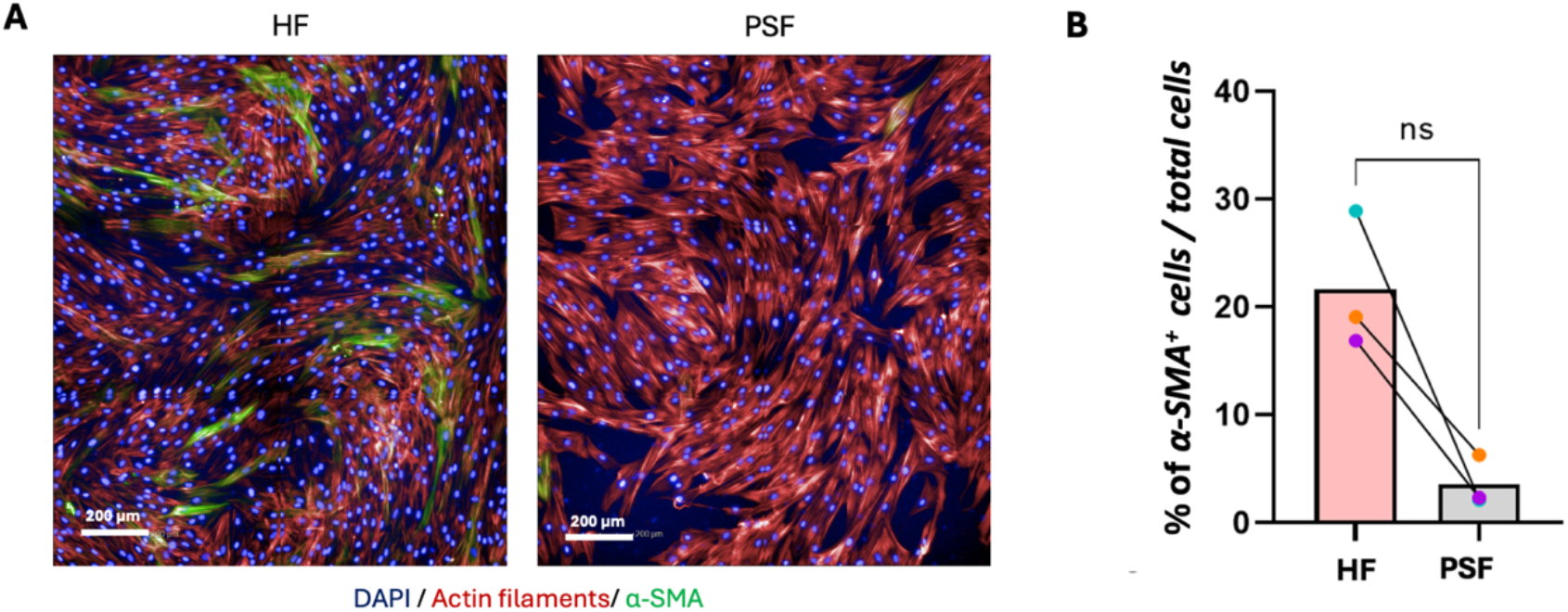
Comparison of α-SMA protein expression capacities of HF and PSF. After 72h of culture in the presence of TGF-β, cells were fixed in PFA and labelled with anti-α-SMA-Alexa Fluor 488 antibody (green) and phalloidin-Alexa fluor 594 (actin, red). The nuclei are labelled with DAPI (blue) and the fluorescence of the cells is analyzed on a high-throughput imaging system. A) Representative images of myofibroblast differentiation from HF (left) and PSF (right). Scale bar 200 µm. B) Quantification of the percentage of α-SMA-positive cells relative to the number of total cells in HF and PSF. n= 3 independent paired samples. Each point represents a different patient sample; non-parametric paired t-test (ns: not significant).

## 3. Discussion

Pressure sores are ischaemic tissue lesions caused by prolonged pressure against a bony surface.The resulting ischaemia affects tissues ranging from the skin to subcutaneous tissues, muscles and fascia simultaneously [1,2]. The development of skin necrosis in a pressure ulcer is therefore the result of tissue hypoxia and ischaemia-reperfusion injury, leading to deep skin ulceration. In spinal cord injured patients, pressure sores represent a major public health problem. These patients suffer from problems related to long treatments, septic risk, and secondary tumorigenesis. Conventional therapies are often insufficient and patients require debridement surgery and flap coverage that may be defective, because of lack of tissue to mobilize [18]. Therefore, sometimes there is no longer any surgical solution.

In this context, fundamental researches on the healing process of pressure sores must be enriched in order to understand any novel therapies that may be applied.

Thus, the study of the cellular behavior of tissues affected by pressure ulcers at baseline could help to understand therapies that may be promising for pressure ulcers in the future.

Fibroblasts from pressure ulcers show a different phenotypic and genotypic profile compared to fibroblasts from healthy skin.

Histological analysis on pressure sores shows significant changes in pressure sores in the architecture of the papillary and reticular dermis with the presence of focal necrosis, macrophage infiltrates and an important fibrosis. Morphological differences were observed as well as a lower fibroblast density in the healthy group compared with the pathological group in relation with fibroscarring.

To compare the genotype of the two types of fibroblasts, we studied the expression of genes coding for matrix molecules, genes coding for proteins involved in the remodeling of the extracellular matrix, as well as genes coding for different cytokines and growth factors. Derma fibroblasts perform a crucial role in wound healing by populating the wound site to produce ECM which forms a scar [19] and several studies show that healthy dermal fibroblasts are naturally selected for wound repair because of their intrinsic ability to produce thick and organized collagen in comparison to wounds with poorly organized ECM [19-24]. Pressure sore fibroblasts express less genes coding for ECM proteins such as COL3A1 and FN1 than healthy fibroblasts and present a reduced expression of genes coding for proteins involved in tissue degradation and remodeling such as MMP3 and TIMP2.

TIMP2 is an inhibitor of several metalloproteinases such as MMP1, MMP2 and MMP3 which have immunosuppressive properties thus promoting skin repair [25]. Activated MMP3 is capable of degrading many ECM components, such as proteoglycans, fibronectins and collagens (III, IV and V) and is mainly inhibited by TIMP1 and TIMP2 [26]. Surprisingly, a study showed that human dermal fibroblasts exposed to wound exudate from chronic venous ulcers overexpress MMP1 and MMP3 while the expression of TIMP1 is decreased [27], and this depends on the stage of the ulcer as well as the wound microenvironment. Our results also show that pathological fibroblasts express less the gene coding for collagen III, which is highly expressed in the early phases of skin repair. It has been shown that the application of mechanical pressure on fibroblasts isolated from a hypertrophic wound also decreases the expression of Col3A1 [28] supporting our results.

The study of the gene coding for the Connective tissue growth factor (CTGF) was equally performed showing a lesser expression in pathological pressure sores fibroblasts. The CTGF is an established fibrotic biomarker and a well-known mediator of inflammatory effects [29]. CTGF signaling activation is crucial for the inflammatory and remodeling process underlying wound repair. It stimulates fibroblasts to release elevated quantities of α-smooth muscle actin (α-SMA), which enhances the contractile potential of these fibroblasts. This could equally be in relation with a lower expression of ACTA2 and a lesser miofibroblastic differentiation in pressure sores fibroblasts’. Fibroblast migration to the wound is regulated by inflammatory mediators such as C5a, fibronectin, PDGF, fibroblast growth factor (FGF), and TGF β [19]. Gene expression studies show that fibroblasts from the pressure sore underexpress the ACTA2 gene, coding for the α-SMA protein, which is confirmed by a lower expression of the α-SMA protein during myofibroblastic differentiation under stimulation of TGF β compared to healthy fibroblasts.

The development of a pressure sore is often due to the application of intense pressure on the tissues inducing ischemia and tissular hypoxia. However, a study has shown that the application of mechanical pressure on fibroblasts isolated from a hypertrophic wound induces a decrease in the expression of the ACTA2 gene associated with dedifferentiation of myofibroblasts [30]. Similarly, culturing fibroblasts from the tail of mice under hypoxia conditions inhibits their ability to differentiate into myofibroblasts [31]. Thus, fibroblasts located in the pressure sore, exposed to pressure forces and hypoxia, lose their ability to differentiate into myofibroblasts, thus confirming our results. However, myofibroblasts play a major role in the closure of the skin lesion thanks to their contraction capacity and this loss of function could partially explain the chronicization of these wounds. However, pathological fibroblasts express more genes coding for pro-inflammatory cytokines such as IL-1β but also anti-inflammatory ones such as IL-10. The overexpression of IL-1β is confirmed by a study comparing mRNA expression in whole pressure ulcer tissue and healthy skin tissue from different patients [32]. This study shows a higher level of mRNA coding for IL-1β in pressure ulcers. However, it has been also showed an increased level of mRNA coding for TNF-α, which is not the case of our results. However, the authors analyze gene expression in the whole pressure ulcer tissue, which contains other cell types, such as macrophages, which may explain this difference. Further analysis of gene and protein expression of different pro- and anti-inflammatory cytokines will be necessary to confirm our results.

Surprisingly, we did not find differences for the hypoxia factor HIF-1α gene expression. As mentioned previously, the formation of a pressure ulcer is induced by a condition of tissue hypoxia. Thus, we hypothesized that the hypoxia factor HIF-1α could be overexpressed in fibroblasts isolated from pressure ulcers. However, our results do not show any difference in expression between the two populations of fibroblasts. This can be explained by the fact that the fibroblasts from the “pressure ulcer” area are extracted from their environment and cultured without hypoxia conditions. This observation allows us to draw a first limitation of this project: fibroblasts are cultured in conditions different from their original environment. Therefore, it would be interesting to culture them in hypoxic conditions as closer to their initial environment and to analyze the same parameters observed in this study.

In our study and as expected, pathological fibroblasts appear to proliferate less quickly than healthy fibroblasts. These results confirm previous studies which have shown that the doubling time of fibroblasts from the pressure sore was greater than that of fibroblasts from the periphery of the ulcer or from the healthy skin [33]. This limit in the proliferation capacity of fibroblasts from the pressure sore is associated with replicative senescence which is detected in tissues but also in cultured fibroblasts [34]. It would therefore be interesting to determine whether our difference in proliferative capacity is also associated with an early entry of pathological fibroblasts into senescence.

All of our results highlight a morphological, genetical and functional difference between healthy and pathological fibroblasts which have a modified phenotype, less effective for skin repair. This suggests that new therapies for chronic wounds must take into account the environment in which they are applied and that pathological cells do not necessarily respond to treatments in the same way as healthy cells. Our results are not statistically significant, although several trends emerge. This is explained by the heterogeneity of the patients’ medical history and requires repetition of the experiments

## 10. Conclusion

In this study, fibroblasts from a pressure sore and from healthy skin were isolated in order to compare their phenotypical and functional characteristics. We showed that pressure sore fibroblasts have a morphological, genetical and functional different profile, and pathological fibroblasts have a reduced proliferation and a reduced myofibroblastic differentiation compared to healthy fibroblasts. However, the migration capacities of the two populations appear to be identical. We also demonstrated that fibroblasts from the pressure ulcer area have decreased expression of genes encoding ECM molecules and proteins involved in tissue remodeling. Overall, our results show that fibroblasts present in the pressure ulcer display an altered phenotype, less favorable for effective skin repair.

However, these experiments were conducted on a limited number of samples; therefore, it will be necessary to increase the number of samples to confirm these results.

